## Supplementary Files for "A nanobody suite for yeast scaffold nucleoporins provides details of the Nuclear Pore Complex structure"

[1]Department of Biology, Massachusetts Institute of Technology, Cambridge, Massachusetts, United States
[2]Boston Children's Hospital and Harvard Medical School, Boston, Massachusetts, United States
*corresponding author

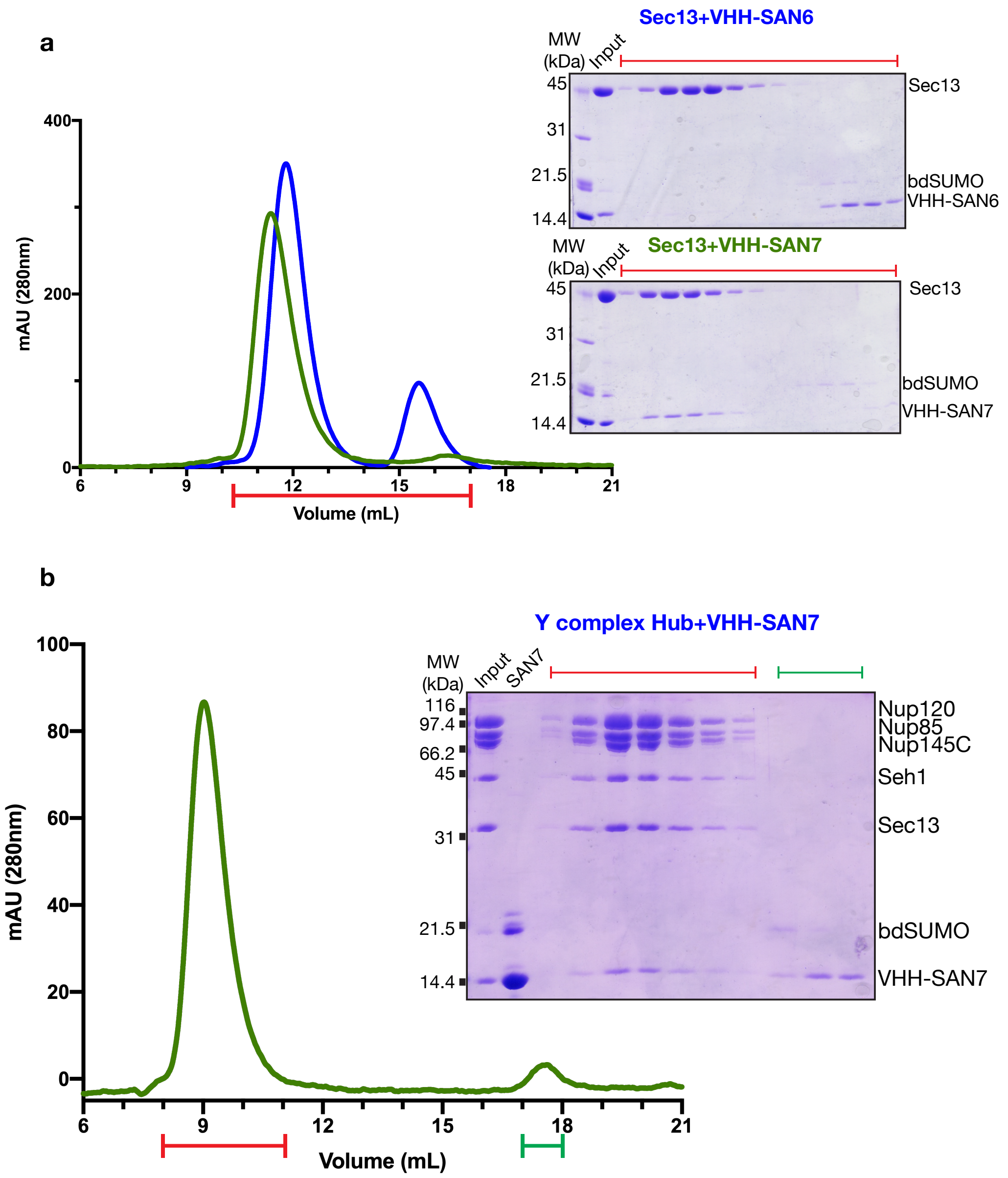


### Supplementary Figure 1.

**VHH-SAN7 binds Sec13 in the Y complex hub**

(a) Size exclusion chromatography (SEC) of Sec13-Nup145C$_{\text{blade}}$ pre-incubated with VHH-SAN6/7 (1:1 molar ratio). SDS-PAGE analysis of SEC experiments for the fractions indicated. Co-migration of Sec13 and VHH-SAN7 indicates complex formation, whereas VHH-SAN6 migrates as a separate peak. (b) SEC with pre-incubated Y complex hub and VHH-SAN7 (1:2 molar ratio). SDS-PAGE analysis of the SEC experiment for the fractions indicated. Co-migration of the hub and VHH-SAN7 shows complex formation.

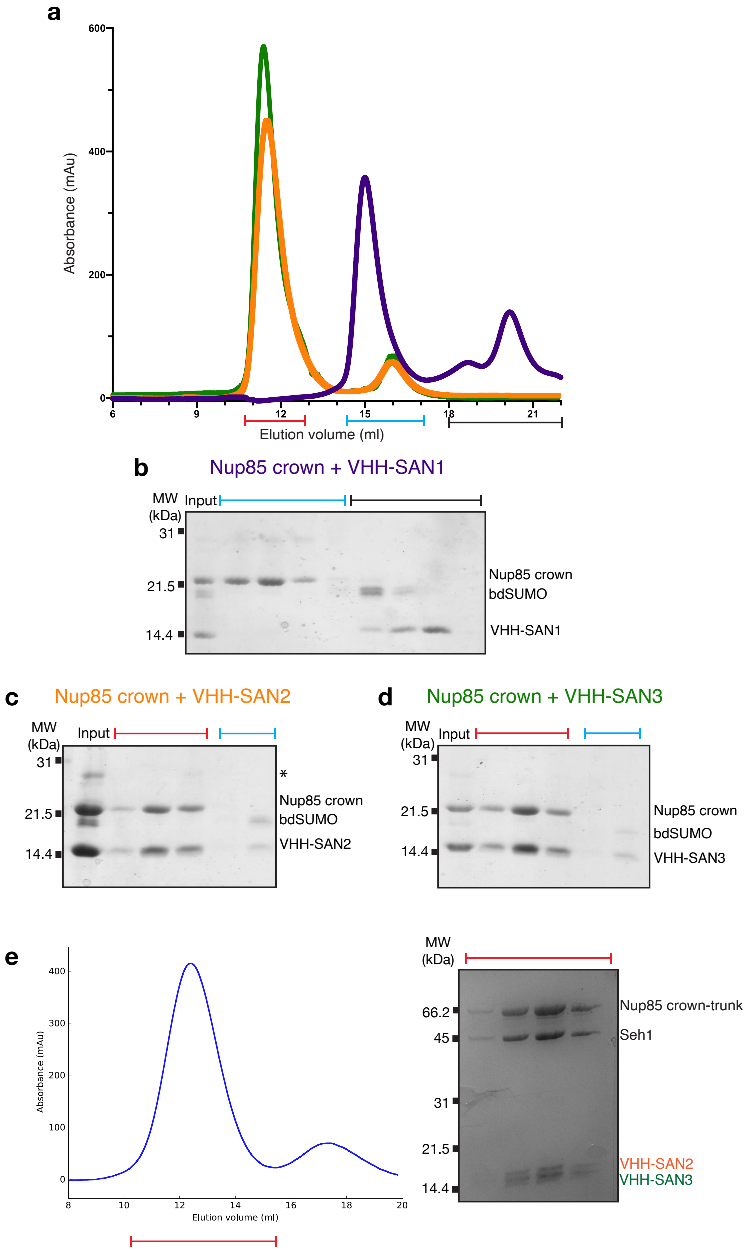


### Supplementary Figure 2.

**VHH-SAN2 and VHH-SAN3 bind the Nup85 crown module**

(a) Size exclusion chromatography (SEC) of Nup85$_{crown}$ pre-incubated with VHH-SAN1 (purple) VHH-SAN2 (orange) VHH-SAN3 (green) (1:1.5 molar ratio), absorbance at 280 nm. (b, c, d) SDS-PAGE analysis of SEC experiments of the indicated fractions for (b) VHH-SAN1 (c) VHH-SAN2 (c) VHH-SAN3. * indicates an unknown contaminant. (e) SEC of Nup85$_{crown-trunk}$-Seh1 pre-incubated with VHH-SAN2/3 (1:1.5 molar ratio), absorbance at 280 nm. Left panel shows SDS-PAGE analysis of the indicated fractions.

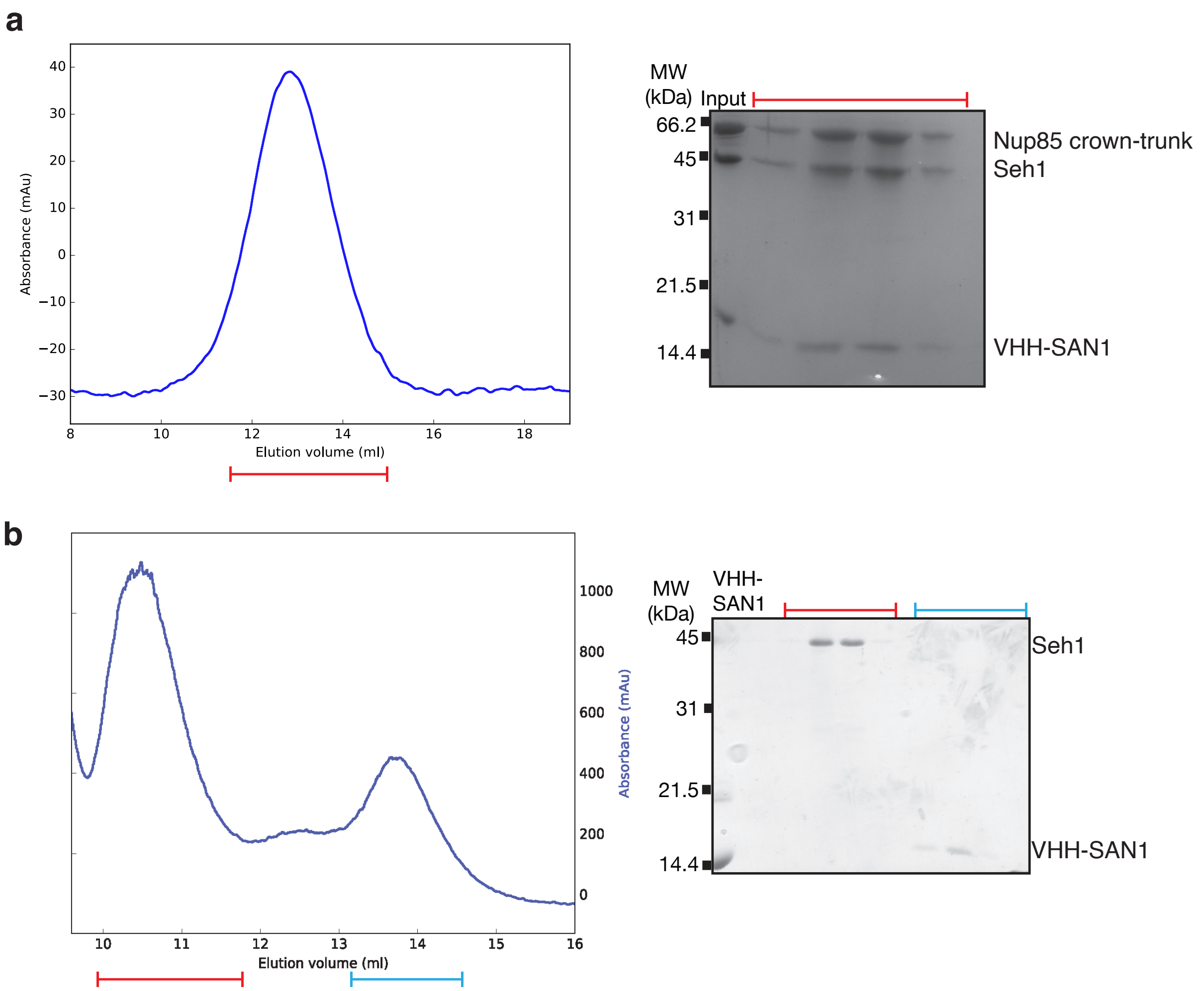


### Supplementary Figure 3.

**VHH-SAN1 binds the Nup85 trunk module**

(a) Size exclusion chromatography (SEC) of Nup85$_{\text{crown-trunk}}$-Seh1 pre-incubated with VHH-SAN1 (1:1 molar ratio), absorbance at 280 nm. Left panel shows SDS-PAGE analysis of the indicated fractions. (b) SEC of Seh1 pre-incubated with VHH-SAN1 (1:1 molar ratio), absorbance at 220 nm. Left panel shows SDS-PAGE analysis of the indicated fractions.

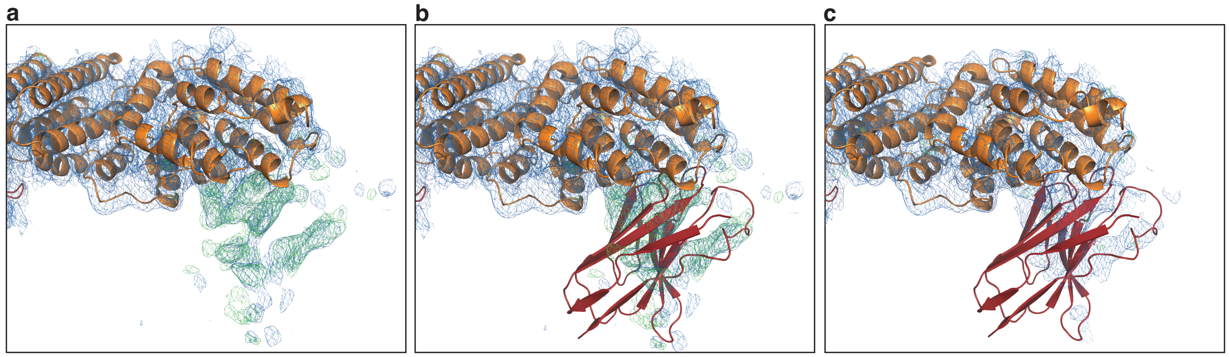


#### Supplementary Figure 4.

**Difference density at the Nup85 crown**

Electron density for the Nup85$_{crown}$ and VHH-SAN2. $2F_O$-$F_C$ map contoured at $1\sigma$ is shown in blue, the $F_O$-$F_C$ map contoured to $2.5\sigma$ shown in green. (a) Density after placing only Nup85-Seh1. (b) Rigid body placement of a VHH-SAN2 model. (c) Final density after refinement with VHH-SAN2. Residues outside of the density for VHH-SAN2 have an occupancy set to zero.

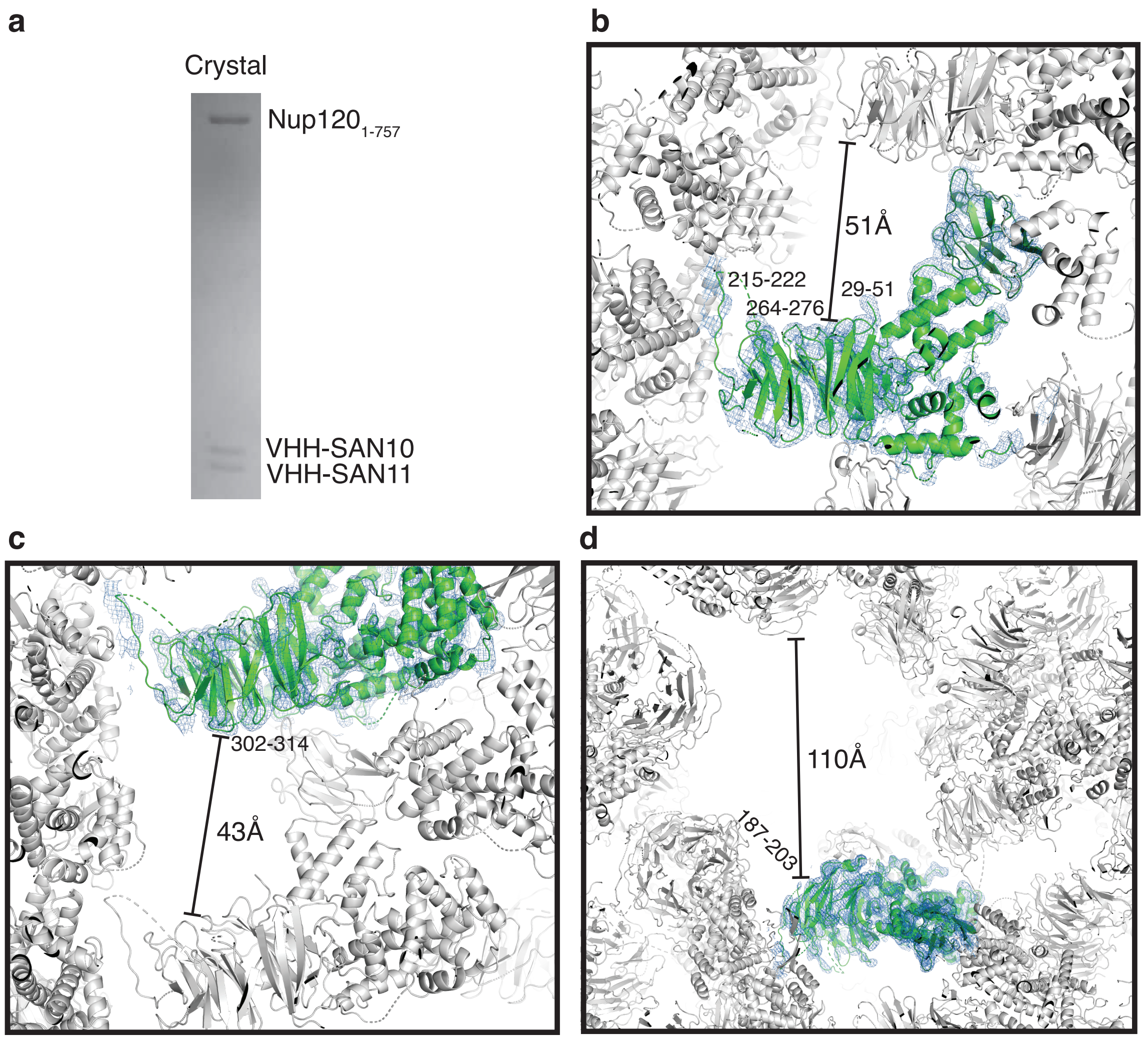


### Supplementary Figure 5.

**Crystal packing in Nup120-VHH-SAN10/11**

(a) SDS-PAGE of $Nup120_{1\text{-}757}$-VHH-SAN10/11. Both nanobodies are present in the crystals. (b, c, d) Views of crystal packing contacts and solvent channels. One copy of $Nup120_{1\text{-}757}$-VHH-SAN11 shown in green with $2F_O\text{-}F_C$ density contoured to $1\sigma$. Symmetry related molecules are shown in gray. Unbuilt loops lacking electron density are listed and distances between molecules are indicated.
